# Sampling from commercial vessel routes can capture marine biodiversity distributions effectively

**DOI:** 10.1101/2022.06.23.497327

**Authors:** Elizabeth Boyse, Maria Beger, Elena Valsecchi, Simon Goodman

## Abstract

Collecting fine-scale occurrence data for marine species across large spatial scales is logistically challenging, but is important to determine species distributions and for conservation planning. Inaccurate descriptions of species ranges could result in designating protected areas with inappropriate locations or boundaries. Optimising sampling strategies therefore is a priority for scaling up survey approaches using tools such as environmental DNA (eDNA) to capture species distributions. eDNA can detect diverse taxa simultaneously, but to date has rarely been applied across large spatial scales relevant for conservation planning. In a marine context, commercial vessels, such as ferries, could provide sampling platforms allowing access to under-sampled areas and repeatable sampling over time to track community changes. However, sample collection from commercial vessels could be biased and may not represent biological and environmental variability. Here, we evaluate whether sampling along Mediterranean ferry routes can yield unbiased biodiversity survey outcomes, based on perfect knowledge from a stacked species distribution model (SSDM) of marine megafauna. Simulations were carried out representing different sampling strategies (random vs systematic), frames (ferry routes vs unconstrained) and number of sampling points. SSDMs were remade from different sampling simulations and compared to the ‘perfect knowledge’ SSDM to quantify the bias associated with different sampling strategies. Ferry routes detected more species and were able to recover known patterns in species richness at smaller sample sizes better than unconstrained sampling points. However, to minimise potential bias, ferry routes should be chosen to cover the variability in species composition and its environmental predictors in the SSDMs. The workflow presented here can be used to design effective eDNA sampling strategies using commercial vessel routes globally. This approach has potential to provide a cost-effective method to access remote oceanic areas on a regular basis, and can recover meaningful data on spatiotemporal biodiversity patterns.

## Introduction

Knowledge of species’ ranges is essential for assessments of conservation status, to detect changes in distributions, and to inform spatial planning decisions (Wetzel et al., 2018). Initiatives to aggregate biodiversity data, including the Global Biodiversity Information Facility (GBIF) and the Ocean Biodiversity Information System (OBIS), have increased access to global standardised datasets (Grassle, 2000; Telenius, 2011). However, these datasets are limited by data quality issues, such as positional accuracy or duplicates of records, and spatial, temporal and taxonomic biases (Moudrý and Devillers, 2020). Marine species and habitats are underrepresented due to the monetary and logistical challenges of collecting data, with up to 50 % of records for marine taxa being collected from coastal regions, or are classified as Data Deficient in IUCN Red List assessments (Dulvy et al., 2014; Hughes et al., 2021). Data limitations increase uncertainty in marine spatial planning prioritisations and could lead to less efficient marine reserve systems (Foley et al., 2010; Bani et al., 2020). Novel methods that provide unbiased data are needed for remote areas to improve our knowledge of species distributions, and their conservation. This paper presents a novel framework to design sampling strategies from commercial vessels, to scale up surveys employing tools such as environmental DNA (eDNA) to record species communities more accurately and comprehensively.

In biodiversity surveys, it is usually infeasible to collect samples at very high coverage across large geographical scales, so sampling strategies target the collection of non-biased data at resolutions relevant to study aims. Design-based sampling methods, including random, systematic, and stratified random sampling, ensure that every sampling unit has a non-zero probability of being sampled (Wang, J.-F. et al., 2012). Model-based sampling designs aim to avoid bias by considering spatial autocorrelation and heterogeneity in the sampling frame, the area to which sampling is restricted (Zhang, C. et al., 2020). The choice of sampling design is dependent on the study objectives and study area characteristics as no method consistently outperforms other methods (Zhang, C. et al., 2020). These sampling designs assume that it possible to access the entire sampling frame for sample collection. However, in the marine environment this is often impossible to achieve, especially when considering the large spatial scales relevant for marine spatial planning, or the conservation of highly mobile species (Di Sciara et al., 2016).

Commercial vessels, such as ferries, typically follow specific shipping routes covering large spatial scales comprehensively, making them effective platforms for replicable sampling transects. Ferry-based sampling is a similar concept to collecting samples close to road networks, which is commonly employed in terrestrial biodiversity surveys due to greater accessibility (Kadmon et al., 2004). The data collected can be biased because the presence of roads directly affects species distributions or because they do not represent the environmental gradients in the whole sampling frame (Kadmon et al., 2004). We therefore need to explore sampling methods which can best capture variability in species distributions from restricted sampling frames, as these often offer us low-cost sampling and accessibility to hard-to-reach areas. Samples from a restricted area (i.e. road networks or ferry routes) can still produce species distribution model predictions similar to samples collected from an unconstrained area if the environmental gradients are adequately captured (Tessarolo et al., 2014). Despite ferries being frequently utilised for visual surveys of cetaceans, there is no precedent for selecting networks of ferry routes to infer species composition in areas not directly covered by the routes (Arcangeli et al., 2017). Emerging survey technologies, such as eDNA sample collection, require multiple point sample locations along individual ferry routes, as well as selecting along which routes to sample (Valsecchi et al., 2021). Understanding which sampling strategies will reduce the inherent bias of restricted sampling frames will allow us to best utilise these low-cost sampling opportunities.

Species distribution models can serve as sampling backgrounds for simulating sampling strategies (Tessarolo et al., 2014). Individual species distribution models can be summed using probability or binary predictions to create a stacked-species distribution model (SSDM) that predicts species richness (Calabrese et al., 2014). Species distribution models only consider environmental constraints on species distributions which can lead to overprediction of species richness when combining multiple models, as biotic mechanisms such as dispersal limitations or resource competition are not accounted for (Gavish et al., 2017). However, using stacking methods based on occurrence probabilities instead of thresholding occurrence probabilities leads to SSDMs which predict species richness similarly to macroecological models, whilst also retaining information on individual species (Calabrese et al., 2014; Distler et al., 2015; Grenié et al., 2020). The use of empirical versus simulated communities allows for complex community “organisation” to be included in sampling simulations and can highlight areas of important conservation interest, i.e. rare species distribution ranges or gradients of diversity (Miller, 2014). We can use the outputs from SSDMs as a benchmark to assess sampling biases associated with different sampling strategies (Braunisch and Suchant, 2010).

This study develops a novel approach for assessing the suitability of different sampling strategies to overcome biases associated with spatially constrained sampling platforms, such as commercial vessel routes. Such a strategy could be used to gain high quality data from pelagic areas that are currently under-sampled due to accessibility and monetary constraints (Hughes et al., 2021). Firstly, we quantify the magnitude of bias of a spatially constrained network of ferry routes with different sampling strategies, relative to unconstrained sampling across the Mediterranean Sea. Second, we consider how environmental variability or species composition impact the effectiveness of ferry routes as a sampling frame with different subsets of ferry routes. Finally, we evaluate the impact of taxonomic sampling biases on correctly predicting gradients in biodiversity as these biases are pervasive in sampling methods such as eDNA metabarcoding. We use ferry routes in the Mediterranean Sea, but the workflow could be applied to shipping networks anywhere.

## Methods

### Building Stacked Species Distribution Models

We assembled a SSDM to represent true species distributions based on observational data from online biodiversity repositories and environmental data. An initial literature search identified 171 species of large marine predators (elasmobranchs, mammals, teleost fish and turtles) with known occurrences in the Mediterranean. We defined a predator based on two criteria; maximum length greater than or equal to 100 cm and a trophic level greater than or equal to four as reported in FishBase (https://www.fishbase.se/) or SeaLifeBase (https://www.sealifebase.ca/). Nine species were retained that only satisfied one of the criteria (Supplementary Information). Occurrence records for species were downloaded from GBIF (https://www.gbif.org/), OBIS (https://obis.org/) and EurOBIS (https://www.eurobis.org/) and supplemented by the Medlem and Accobams datasets for elasmobranchs and cetaceans respectively (ACCOBAMS Survey Initiative, 2018; Mancusi et al., 2020). We subset occurrences to include records from 2000 onwards to correspond with the years that environmental variables were available. We removed occurrence records where GPS coordinates had fewer than three decimal places to improve positional accuracy, and duplicates between the datasets based on species, coordinates, year and month (Moudrý and Devillers, 2020). Records which had the same species, year and month but different coordinates as a result of potentially rounding between databases were also assumed to be duplicates and removed manually. After quality checking, we only retained species with 40 or more occurrence records to improve model accuracy, leading to 43 species in the final presence only dataset, with records for individual species ranging from 41 to 7,822 occurrences (Meynard et al., 2019). The selected species were representative of all marine vertebrates including teleost fish (n=20), elasmobranchs (n=13), marine mammals (n=9) and one sea turtle species (Table S1, Supplementary Information) To account for sampling bias in data repositories, occurrences were spatially thinned with a nearest neighbour distance of 10 km using the spThin R package (Aiello-Lammens et al., 2015). This approach prevents clusters of occurrences although does not account for large scale spatial biases. This method resulted in less than 40 occurrences for ten species, in which case the original data was used instead. We downloaded six environmental predictors (bathymetry, sea surface temperature mean, sea surface temperature range, chlorophyll *a* mean, bathymetry slope and distance from shore) from Bio-ORACLE and Marspec in WGS84 projection at a resolution of 0.83×0.83 degrees (Sbrocco and Barber, 2013; Assis et al., 2018). These environmental variables are of known importance to marine predators, or their prey species (Azzellino et al., 2012; Klippel et al., 2016; Lambert et al., 2017). These variables were normalised to between 0 and 1 to account for units differing by orders of magnitude.

We modelled individual species distributions with three different approaches, maximum entropy (MAXENT), multiple adaptive regression splices (MARS) and random forest (RF). MAXENT was run with 10, 000 random background points using the dismo R package (Hijmans et al., 2017). We selected presence-absence algorithms MARS and RF despite having a presence-only dataset as they perform better than presence-only models, when employed with pseudo-absence data (Barbet-Massin et al., 2012; Zhang, L. et al., 2019). We generated 1000 pseudo absences for MARS and an equal number of pseudo absences as presences for RF, both randomly selected within a restricted sampling frame using the two-degree method as recommended by Barbet-Massin et al. (2012). We allowed first order interactions to be fitted for MARS (Wisz et al., 2008). RF was run with 5000 regression trees and a terminal node of 5 (Zhang, L. et al., 2019). We randomly assigned the data set into training (70 %) and testing (30 %) sets three times for cross validation (Sundaram and Leslie, 2021; Arenas-Castro et al., 2022). We assembled the model projections across the three modelling methods using weighted AUC scores for each species. Probabilities of occurrence were translated to binary occurrences using the sensitivity (i.e. true positive rate) equals specificity (i.e. true negative rate) threshold (Liu et al., 2005). The species binary ensemble models were then summed to create the final SSDM showing species richness (Figure 1). Binary SSDMs were selected as binary data was required for sampling simulations. This initial SSDM created with occurrence data from online repositories will be referred to as the ‘perfect knowledge’ SSDM for sampling simulations comparisons. All species distribution modelling was carried out using the SSDM R Package using R version 4.1.0 (Schmitt et al., 2017; R Core Team, 2021).

**Figure 1.**
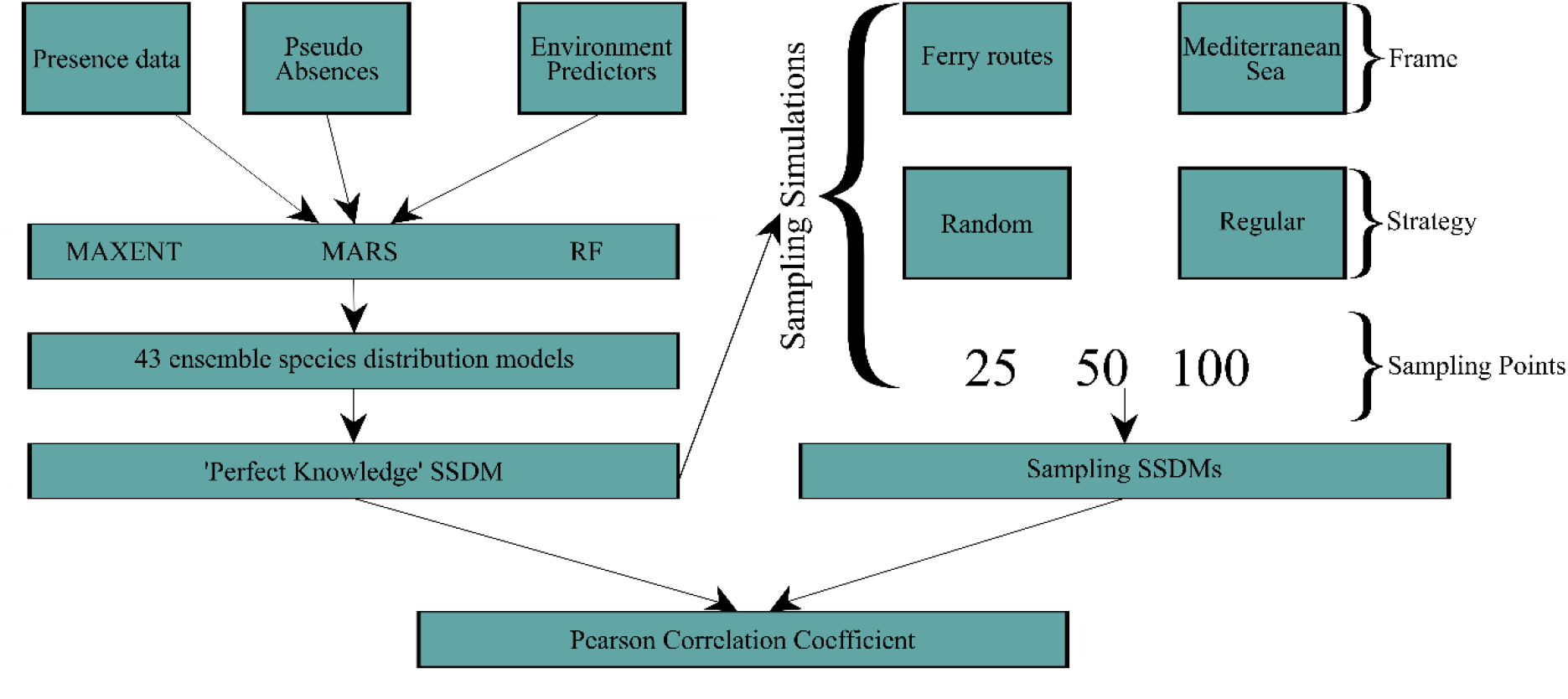
Schematic diagram showing the workflow to create the ‘perfect knowledge’ SSDM using occurrence data from online repositories and extracting occurrence data from the ‘perfect knowledge’ SSDM to build the sampling SSDMs. Sampling SSDMs were compared to the ‘perfect knowledge SSDM to evaluate their predictive capacity.

### Sampling strategy simulations

To enable comparisons of different sampling strategies relative to the ‘perfect knowledge’ SSDM, we selected fifteen operational ferry routes of varying lengths (both intra/inter country tracks) to represent the distribution of ferry routes in the Mediterranean (Figure 2A). We simulated two sampling strategies (random and regular) across different sample sizes (25, 50, 100 sampling points) with either the ferry route network or the Mediterranean as a sampling frame to compare differences between biodiversity detected by a restricted sampling frame versus unconstrained sampling (Figure 1). We explored different combinations of ferry routes, referred to as ‘ferry subnetworks’, to consider the importance of environmental and species data coverage by the ferry routes. We simulated each sampling strategy combination 1000 times to calculate the mean number of species sampled, and the mean number of occurrences per species in the simulations. All sampling simulations were carried out using the spsample() function from the sp R package (Bivand et al., 2008).

**Figure 2.**
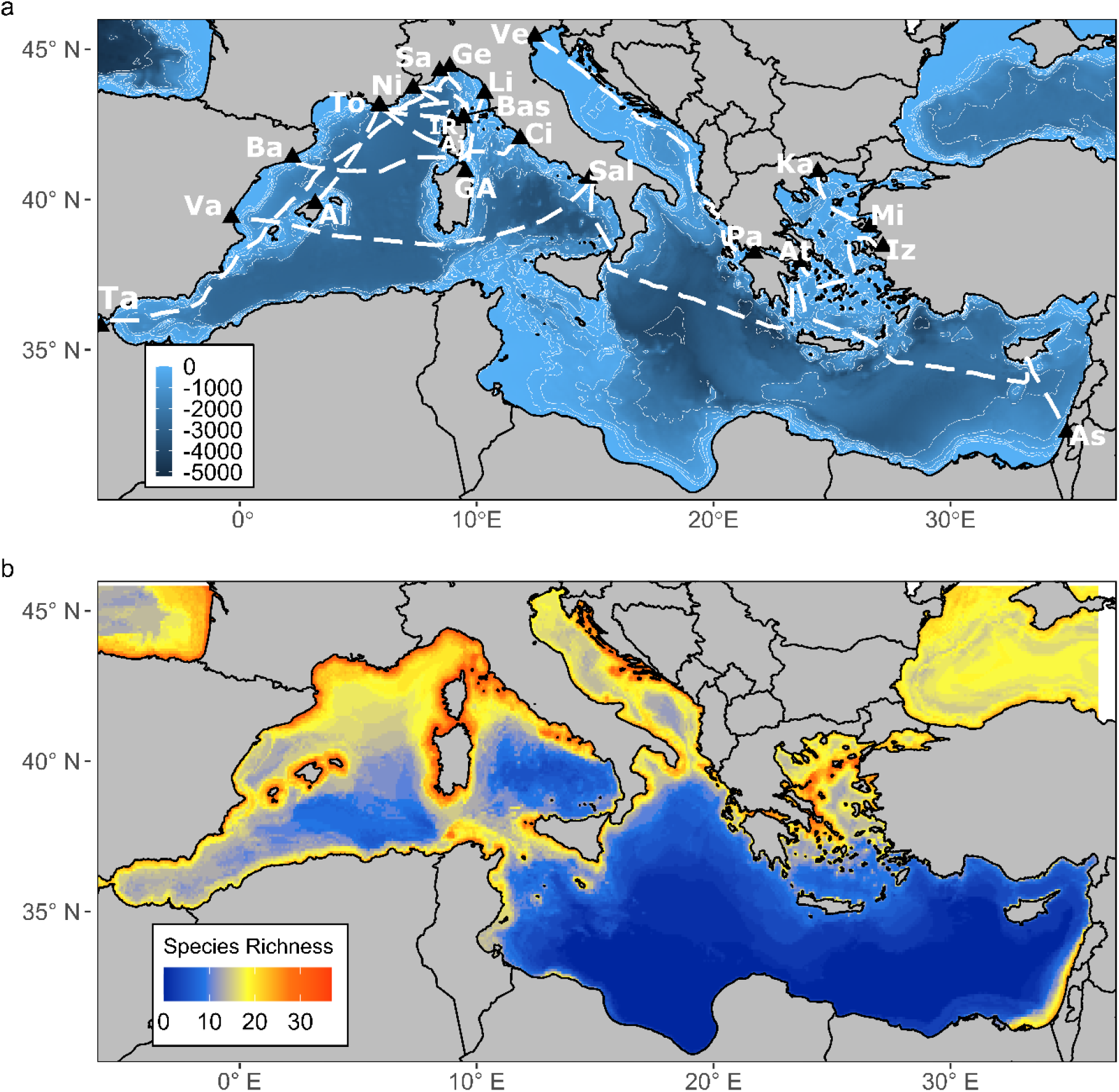
a) Map showing the layout of the whole ferry route network consisting of 15 individual ferry routes. Abbreviations for Ports: Al=Alcudia, Aj=Ajaccio, As=Ashdod, At=Athens, Ba=Barcelona, Bas=Bastia, Ci=Civitavecchia, GA=Golfo Aranci, Ge=Genoa, IR=Ille Rousse, Iz=Izmir, Ka=Kavala, Li=Livorno, Mi=Mitilini, Ni=Nice, Pa=Patras, Sa=Savona, Sal=Salerno, Ta=Tangier, To=Toulon, Va=Valencia, Ve=Venice. b) Original binary stacked species distribution model of 43 marine predators in the Mediterranean using occurrence data obtained from online repositories.

For each sampling strategy simulation, species occurrences were extracted from the ‘perfect knowledge’ SSDM to regenerate new SSDMs from the simulated sampling data, referred to as ‘sampling SSDMs’. We created these SSDMs as before, except that species were not subsampled prior to modelling and species with >20 occurrences were retained. We chose this threshold to evaluate the effect of small sample sizes on model prediction accuracy. To compare species richness across the Mediterranean and ferry route network as sampling frames, 40 replicate SSDMs were built for each combination of sampling frame, size and strategy, using 40 different sampling simulations. We assessed the correlation between the ‘perfect knowledge’ SSDM and the sampling SSDMs based on Pearson’s correlation coefficient to evaluate the effectiveness of different sampling strategies. A three-way Analysis of Variance (ANOVA) was performed to evaluate the effects of sampling strategy, frame and number of sampling points on the correlation coefficient.

### Ferry route subnetworks

We built different ferry subnetworks to evaluate how different coverage of environmental variability and community composition affected the predictive capacity of ferry routes as a sampling frame. The environmental predictors were collapsed into a single index of environmental variability using principal component analysis to quantify the main gradients of environmental variability in the study area (Supplementary Information). The first four principal components explained >80 % of the variability in the environmental predictors. Therefore, we collapsed these principal components by summing the site scores of each principal component weighted according to its contribution (Long and Fisher, 2006; Maina et al., 2008). The resulting environmental variability map was normalised between zero and one, where zero and one represent the most different environments. We quantified climatic bias for different ferry subnetworks by comparing the difference in density functions between environmental variability over the whole study area and those covered by the ferry routes. We split the density functions into 5 equal bins of 0.2 to calculate the climatic bias index. We define our climatic bias index as the sum of the differences of density functions of environmental variability. Salerno-Ashdod was the only ferry route that covered the eastern basin and environmental variability between 0-0.2. Venice-Patras was the only ferry route encompassing the Adriatic Sea and environmental variability 0.6-1. These two ferry routes were therefore used to create the environmental subnetwork as they covered all environmental variability in the study area (Figure3C).

We also considered how community composition differed between the ferry routes. For each ferry route, species occurrences were extracted from each grid cell of the ‘perfect knowledge’ SSDM that overlapped with the ferry route. The number of grid cells that a species occurred in per route was treated as an abundance estimate. We applied a Hellinger transformation to the resulting species abundance x ferry route matrix to dampen the inflated abundances from longer ferry routes (Legendre and Gallagher, 2001). This transformed matrix was then used to create a Bray-Curtis dissimilarity matrix and differences in species composition between ferry subnetworks were quantified by Nonmetric Multidimensional Scaling (NMDS). The NMDS analysis confirmed, as expected, that ferry routes closer together had more similar species composition, with the main cluster formed from routes in the north-western basin (Supplementary Information). This cluster was used to create a deliberately biased ferry route subnetwork (Figure3A). We also used the NMDS analysis to reduce the number of ferry routes from the original ferry route network by randomly selecting one ferry route from each cluster on the NMDS plot to create a subnetwork representing community composition. This reduced the number of ferry routes in the original network from 15 to 9 (Figure 3B).

**Figure 3.**
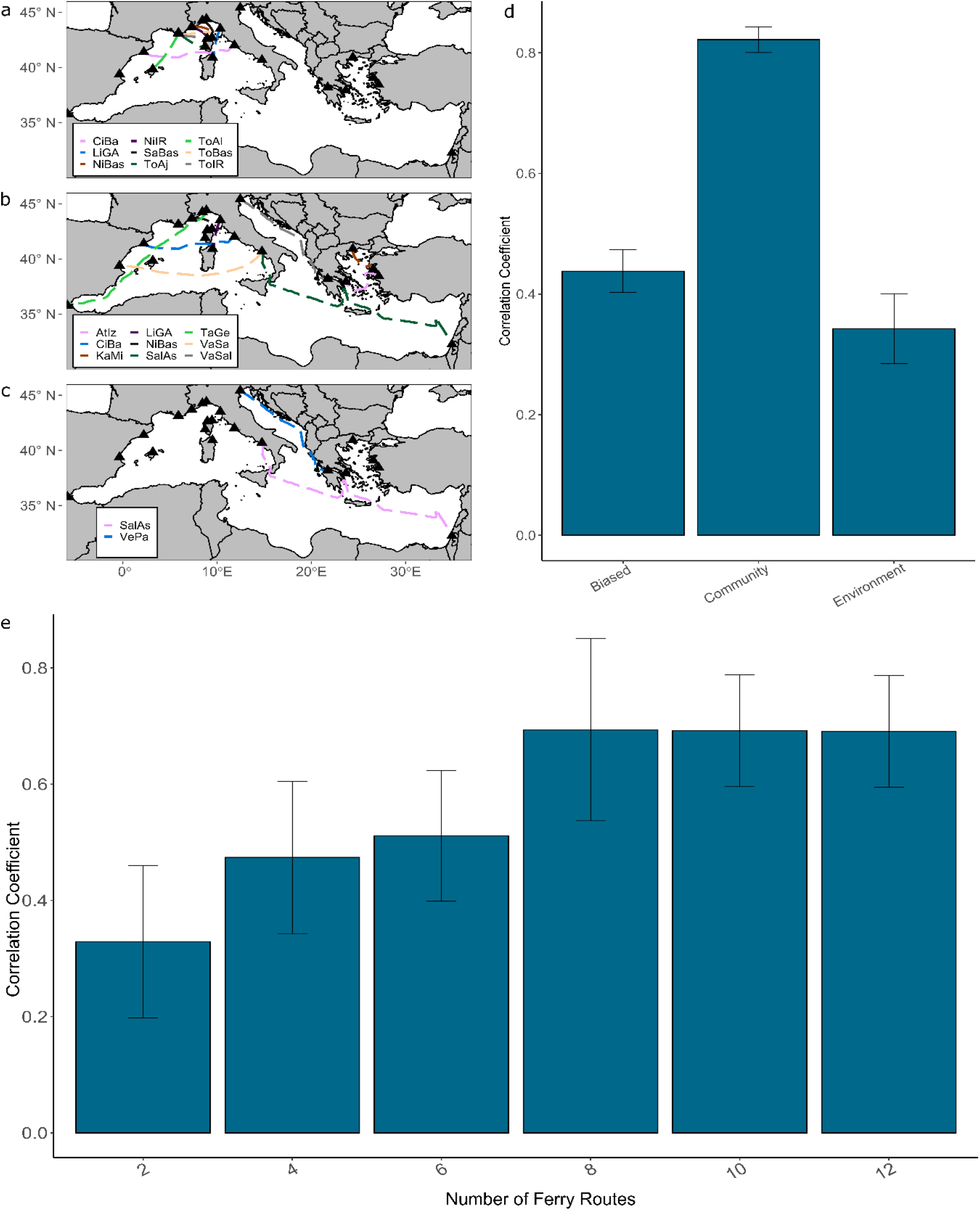
a-c) Maps showing the layouts of the a) biased ferry route subnetwork, b) community ferry route subnetwork, c) environment ferry subnetwork. d) Mean Pearson correlation coefficient between the original S-SDM and sampling S-SDMs for 10 replicate simulations across ferry route subnetworks using 50 regular sampling points. e) Mean Pearson correlation coefficient between original S-SDM and sampling S-SDMs for 10 replicate simulations across subnetworks with differing numbers of ferry routes.

We also produced ferry subnetworks with differing numbers of ferry routes, including 2, 4, 6, 8, 10 and 12 ferry routes by randomly selecting routes from the original ferry route network to evaluate the importance of the number of ferry routes. We built 10 sampling SSDMs using 50 regular sampling points per ferry route subset and compared to the ‘perfect knowledge’ SSDM with Pearson’s correlation coefficient. We assessed the difference between biased, community and environmental subnetworks, and the difference between subnetworks with differing numbers of ferry routes with one-way ANOVAs. We performed post-hoc pairwise comparisons with the Tukey’s Test.

### Taxonomic biases in data collection

The sampling SSDMs were constructed with occurrence data from every sampling point that overlapped with the species distribution. Realistically, no methods for collecting biodiversity data have perfect rates of detectability, so understanding how imperfect detection affects predictions of biodiversity patterns or gradients in biodiversity is important. All biodiversity monitoring techniques, including eDNA metabarcoding, suffer from taxonomic biases (Balint et al., 2018). However, it is unclear how such uncertainty can in turn bias SSDM predictions. To quantify the effect of taxonomic bias, we either removed taxa (Chondrichthyes or Mammalia) or a random subset of species before individual species distribution models were stacked. The random species subset removed the same number of species as the equivalent taxonomically biased model. The models were then compared to the ‘perfect knowledge’ SSDM using Pearson’s correlation coefficient. We analysed the effect of removing specific taxa with a three-way ANOVA and post-hoc pairwise comparisons with the Tukey’s Test.

## Results

### Stacked species distribution model

The SSDM of marine predators in the Mediterranean revealed two main gradients in species richness (Figure 2B). There was higher species richness in the north-western basin compared to the south-eastern basin, and higher species richness nearer to shore. The environmental variable with the greatest influence on model predictions was mean sea surface temperature, whilst the variable with the least influence was bathymetric slope (Supplementary Information). The remaining variables, mean bathymetry, mean chlorophyll concentration, mean temperature range and distance from shore, contributed equally to model predictions. The model tended to overpredict species richness although the extent varied greatly (species richness error mean = 19.06 ± 7.23 s.d). The proportion of presences that were correctly predicted (sensitivity = 0.98 ± 0.12 s.d) was much higher than the proportion of absences correctly predicted (specificity = 0.54 ± 0.17 s.d) (Supplementary Information).

### Comparison of Ferry Route Sampling Frame to whole Mediterranean

The number of species with enough occurrences for modelling (>20) was consistently higher for samples collected along the ferry route network compared to unconstrained sampling across the Mediterranean Sea (Figure 4A). For the smallest number of sampling points (25), only the ferry routes could detect any species with enough occurrence points for modelling (random = 6.3 ± 1.27 s.d, regular = 6.17 ± 0.91 s.d). With 50 sampling points, the ferry routes (random = 18.56 ± 1.53, regular = 18.69 ± 1.99 s.d) detected double the amount of species compared with the Mediterranean (random = 9.42 ± 2.37, regular = 9.67 ± 1.74). The sampling strategy, random vs regular, had no effect on the number of species detected in both the ferry route and whole Mediterranean simulated sampling. The number of species detected increased quickly at small sample sizes but asymptotes between 200 and 500 sampling points where only 5 new species were detected using the ferry route network, and 7 species using the Mediterranean.

**Figure 4.**
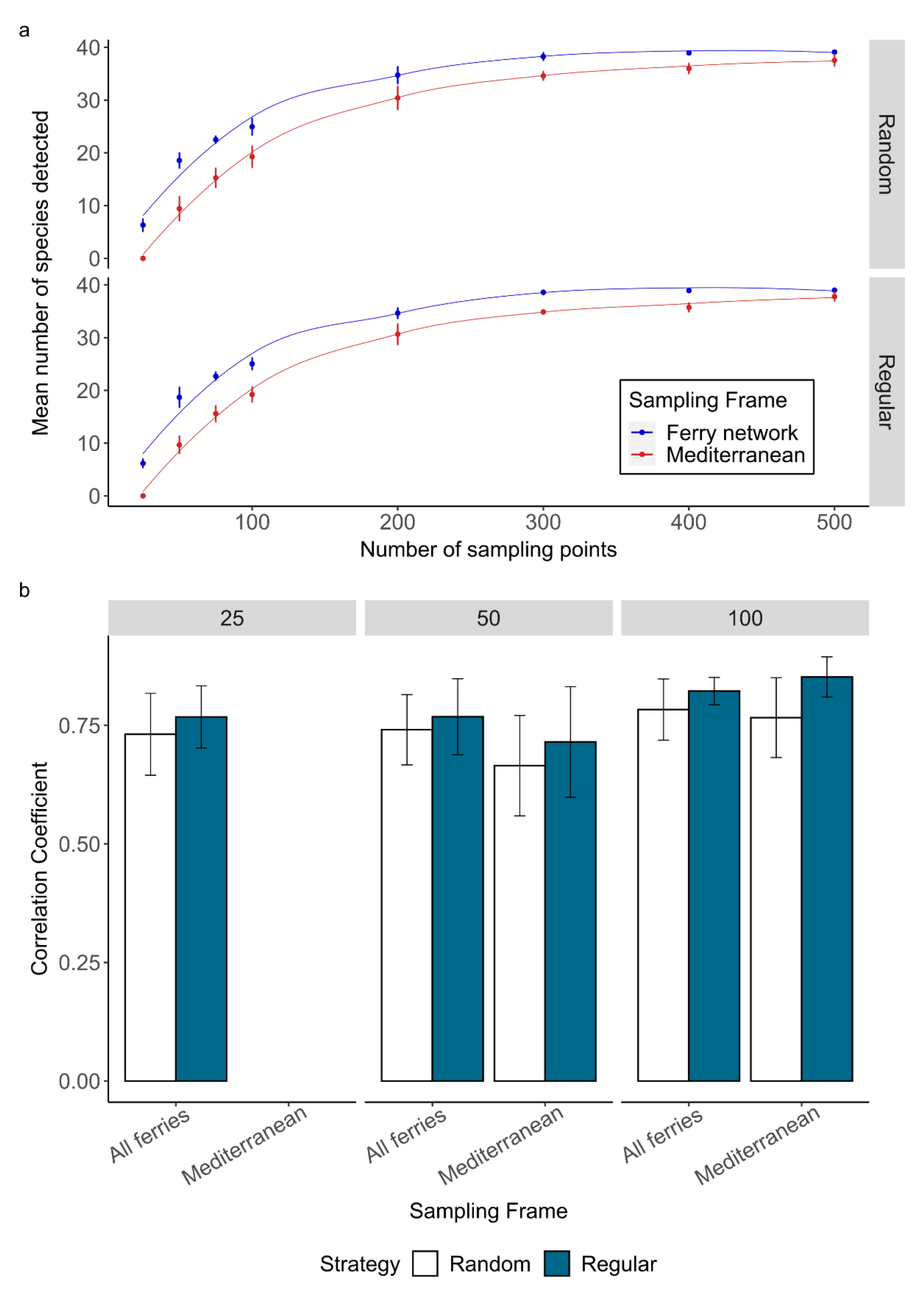
a) The mean number of species detected, with standard deviation bars, across different number of sampling points using either the ferry network or Mediterranean as a sampling frame and either random or regular sampling strategy. b) Mean Pearson correlation coefficient between the original S-SDM and sampling S-SDMs for 40 replicate simulations across the ferry network and Mediterranean for 2 sampling strategies (random and regular) across 3 different sample sizes (25, 50 and 100 sampling points).

Sampling SSDMs with 100 sampling points collected regularly across the Mediterranean were most correlated to the ‘perfect knowledge’ SSDM (85.2 % ± 4 sd) (Figure 4B). Sampling SSDMs produced from 100 sampling points collected either regularly (82.2 % ± 3 sd) or randomly (78.3 % ± 6 sd) across the ferry route network also produced SSDMs highly correlated with the ‘perfect knowledge’ SSDM. Sampling SSDMs produced from the ferry route network differed less between sample sizes than sampling SSDMs from across the Mediterranean (F_(1,373)_=15.8, p <0.001). The sampling SSDM that correlated least with the ‘perfect knowledge’ SSDM was produced from 50 sampling points randomly collected across the Mediterranean (66.5 % ±10). Sampling strategy had a larger effect on the SSDM produced when samples were collected across the Mediterranean compared to the ferry routes (F_(1,373)_=3.91. p = 0.05). A regular sampling strategy produced SSDMs more correlated to the ‘perfect knowledge’ SSDM for 50 sampling points (71.5 % ±12 vs 66.5 % ±10) and 100 sampling points (85.2 % ±4 vs 76.6 % ±8) across the Mediterranean.

**Table 1.** Three-way ANOVA table to evaluate impact of sampling strategy, sampling frame and number of sampling points on correlation coefficients between the sampling SSDMs and the ‘perfect knowledge’ SSDM.

| Factor | Df | Sum Sq | Mean Sq | F value | P value |
| --- | --- | --- | --- | --- | --- |
| Strategy | 1 | 0.4564 | 0.45651 | 37.4637 | <0.001* |
| Size | 2 | 1.1520 | 0.57601 | 47.2701 | <0.001* |
| Sampling frame | 1 | 0.0960 | 0.09603 | 7.8804 | 0.005* |
| Strategy:Size | 2 | 0.0471 | 0.02355 | 1.9324 | 0.15 |
| Strategy:Sampling frame | 1 | 0.0477 | 0.04770 | 3.9147 | 0.049* |
| Size:Sampling frame | 1 | 0.1925 | 0.19249 | 15.7964 | <0.001* |
| Strategy:Size:Sampling frame | 1 | 0.0097 | 0.00966 | 0.7927 | 0.37 |

### Ferry route subnetworks

Different ferry subnetworks varied in their ability to accurately capture community composition in the ‘perfection knowledge’ SSDM (F_(1, 26)=_342.96, p < 0.001) (Figure 3D). The community subnetwork was able to predict the original community composition ∼40 % better than either the biased or environment subnetworks (Tukey’s, p<0.05). The community subnetwork also had a similar climatic bias index to the network with all ferry routes included (Supplementary Information). The environment subnetwork predicted community composition (34.3 % ±0.6) ∼9 % worse than the deliberately biased subnetwork (43.8 % ±0.4) (Tukey’s, p<0.05). The deliberately biased sampling strategy had the highest climatic bias index, whilst the environment subnetwork performed similarly to the original ferry network (Supplementary Information). The number of ferry routes included in a network effected its predictive capacity (F_(1, 54)=_15.286, p<0.001), with correlation to the ‘perfection knowledge’ SSDM increasing from 32.8 % ±13 for networks with 2 ferry routes, to 69.3 % ±16 with 8 ferry routes (Tukey’s, p<0.001)(Figure 3e). Increasing beyond 8 routes does not improve the predictive capacity of the sampling frame but reduces the variability related to which ferry routes are selected in the sub-network (Tukey’s, p>0.05).

### Taxonomic biases in data collection

When species in the same class were stacked together and compared to the ‘perfect knowledge’ SSDM, Actinopterygii (91.9 %) and Chondrichthyes (90.9 %) both had similar species richness patterns to the ‘perfect knowledge’ SSDM (Figure 5). The Mammalia only SSDM (67.96 %) showed weaker correlation with the ‘perfect knowledge’ SSDM. Sampling SSDMs with different taxa removed affected the predicted community composition (F_(1,116)_=8.72, p<0.001)(Figure 6). Sampling SSDMs with Mammalia species removed improved the predictive capacity by 10 % compared to sampling SSDMs with Chondrichthyes removed, or by 7 % compared to random subset of species removed (Tukey’s, p<0.001). This pattern was consistent across a range of sampling sizes and strategies.

**Figure 5.**
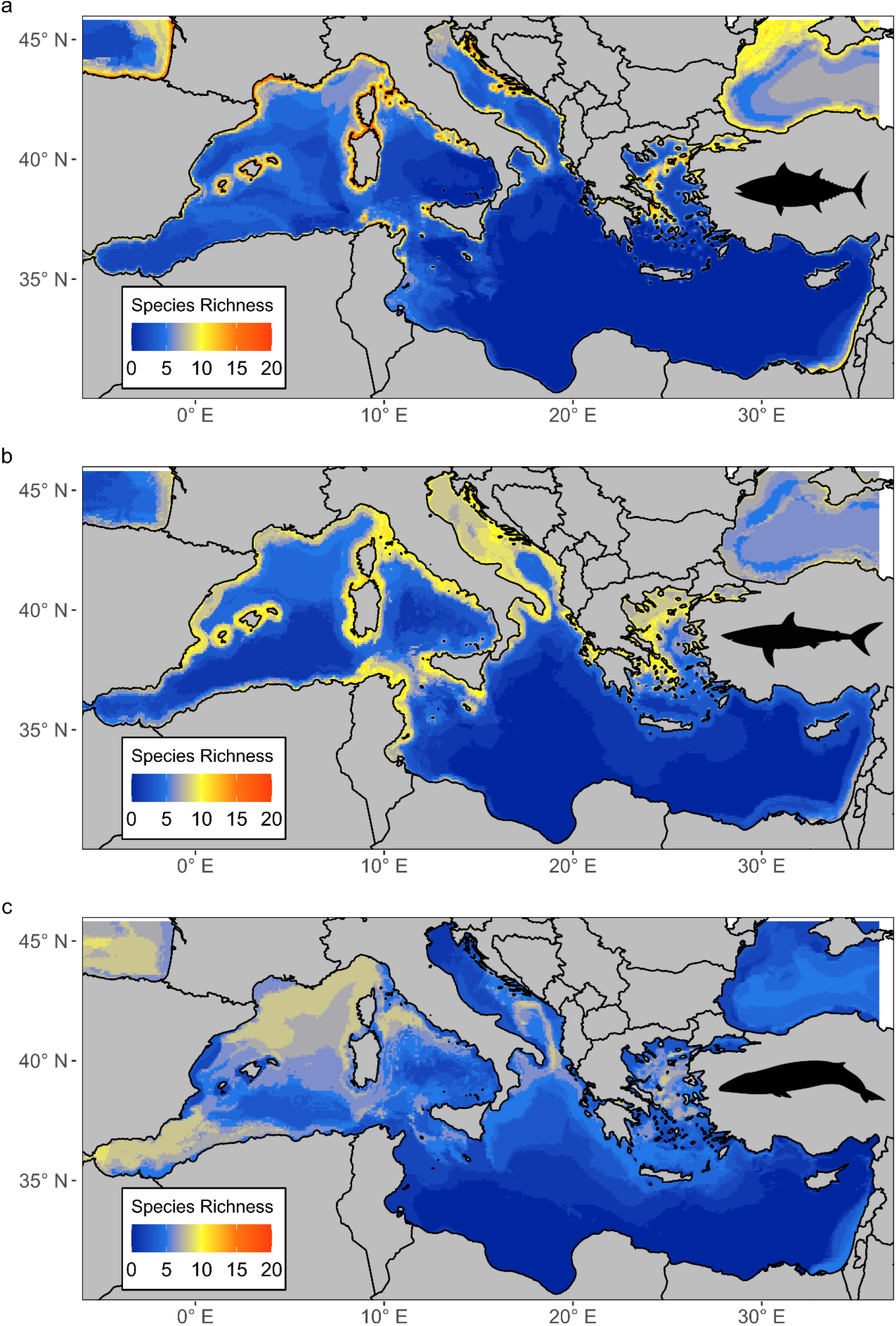
Stacked species distribution models for Class a) Actinopterygii, b) Chondrichthyes, c) Mammalia.

**Figure 6.**
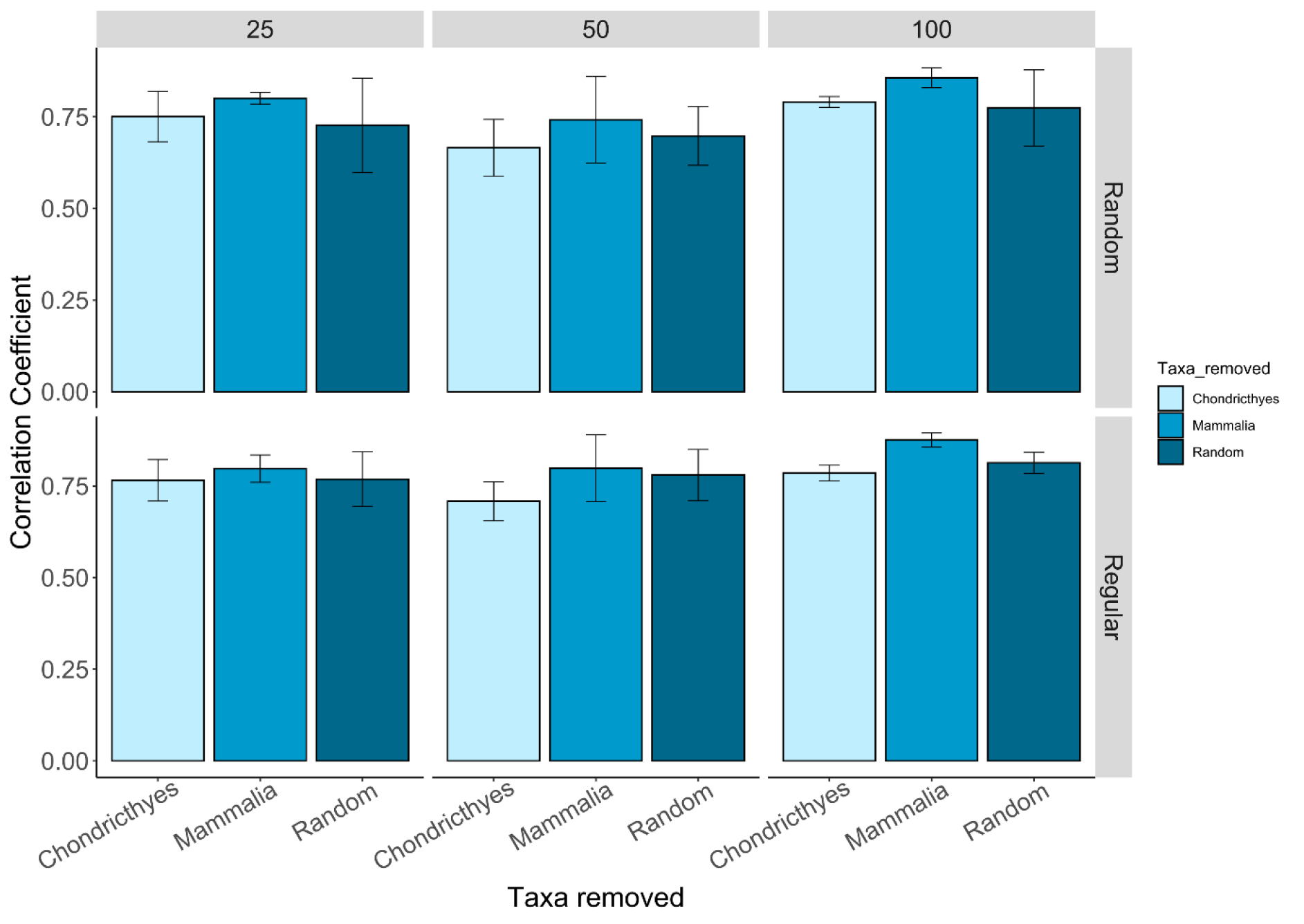
Mean Pearson correlation coefficient between the original S-SDM and sampling S-SDMs with different taxonomic biases for samples collected using the ferry network sampling frame.

## Discussion

Biased sampling remains a key hurdle to predicting biodiversity patterns (Tydecks et al., 2018; Hughes et al., 2021; Moussy et al., 2022). We evaluated the feasibility of using biased sampling frames (in this case commercial vessels) as sampling platforms for collecting species occurrence data for marine species distribution modelling. In this study, we test ferry routes that offer low-cost sampling platforms in hard-to-reach pelagic regions, but introduce biases because spatial sampling is restricted to the routes covered. We found that the inherent bias associated with restricted sampling frames did not lead to a loss in predictive capacity. In fact, for our case study, at small sample sizes (25 sampling points), using ferry routes concentrated sampling to areas with high biodiversity leading to the ferry routes recovering species richness gradients more accurately than unconstrained sampling. This further highlights the cost-effectiveness of ferry routes as sampling platforms and demonstrates that high quality biodiversity data can be recovered from restricted sampling frames. The workflow should be applicable to designing surveys across the global shipping network, including from other vessel types (e.g. container ships), which might serve as future platforms for eDNA sampling for areas that are currently inaccessible on a routine basis (Valsecchi et al., 2021).

### Marine predator SSDM

The SSDM shows that predator species richness is much higher in the north-western basin (Figure 2). This result is unsurprising due to the Strait of Gibraltar linking the western basin to the Atlantic Ocean allowing migration of predators into the Mediterranean (Coll et al., 2010). The north-western basin also has important breeding and foraging habitats for cetaceans which has been acknowledged through the implementation of the Pelagos Sanctuary for Marine Mammals (Notarbartolo-di-Sciara et al., 2008). However, there has been a strong sampling bias in the north-west and coastal regions of the Mediterranean, likely driven by greater economic resources in northern basin countries which benefit from European Union (EU) funding for survey and conservation initiatives (Coll et al., 2010; Amengual and Alvarez-Berastegui, 2018). This bias was reflected in the occurrence data used to create the ‘perfect knowledge’ SSDM, with the density of occurrence points being greatest in the northwestern region and fewest occurrence points furthest offshore in the southern basin. The binary SSDM tended to overpredict species richness, as has been previously reported (Pottier et al., 2013). Combining SSDMs with macroecological constraints may reduce overprediction by accounting for biotic interactions (Guisan and Rahbek, 2011; d’Amen et al., 2015). However, SSDMs can provide similar predictions to macroecological models or joint species distribution models when using a probabilistic stacking approach (Calabrese et al., 2014; Zurell et al., 2020). Despite its limitations, we chose to use a binary stacking procedure as we required presence data to re-run species distribution models from the simulated sampling strategies and the model represents realistic community patterns as a base for sampling simulations.

### Comparison of Ferry Route Sampling Frame to whole Mediterranean

Our selected 15 operational ferry routes are assumed to be representative of the spatial extent of the Mediterranean wide ferry network (Figure 2). Using this ferry route network as a sampling frame achieved species distribution models that predicted the known community from the ‘perfect knowledge’ SSDM as well as or better than samples collected across the whole Mediterranean. Ideally, occurrence data for species distribution modelling would represent a random sample from the population of interest across the entire study area (Araujo and Guisan, 2006). However, geographically biased sampling strategies, i.e. samples only collected close to road networks, can still produce accurate models as long as the environmental predictors are not also biased, as is the case with the ferry route network (Kadmon et al., 2004; Tessarolo et al., 2014). Here, we demonstrate that with smaller sample sizes, samples collected from the biased sampling frame produced more accurate models than samples collected from across the whole Mediterranean Sea (Figure 4). It is more feasible to routinely collect samples on board ferries than to implement dedicated research surveys over large spatial scales comparable to the Mediterranean Sea. Therefore, we show that routine sampling on ferries, such as eDNA sampling, can serve as an important approach to conduct representative biodiversity sampling (Valsecchi et al., 2021). Since fewer samples are required to produce models with similar accuracy from ferry routes compared to the whole Mediterranean, this further increases the cost-effectiveness of ferry routes as a sampling platform. For the ferry route network, there is no cost benefit to doubling the sample size as this does not improve the SSDM community composition prediction (Figure 4). However, the SSDM made with 25 sampling points only detected between 5-8 species (11-18 %) whereas SSDMs with 50 sampling points detected 16-21 (37-48 %) species, and SSDMs with 100 sampling points detected 19-26 species (44-60 %) (Figure 4). If the aim of the study is to look at patterns in species richness, such as gradients in diversity, then a small sample size is adequate. However, if individual species distributions, or the detection of rare species is also important, then larger sample sizes will be required. These sample sizes are based on 100 % detection rate of the species when they are present which is unrealistic for any sampling method. However, we expect that the patterns observed between sample sizes and sampling frames should hold true as long as the detection probabilities are constant across sampling frames. Sampling SSDMs from the ferry networks were less effected by sampling strategy than sampling SSDMs from the whole Mediterranean, where random sampling consistently produced more poorly performing SSDMs. By limiting the available sampling frame to such an extent, this potentially reduces the impact of sampling strategy, and prevents random sampling from forming clusters which do not cover the study area’s environmental variability (Zhang, C. et al., 2020). These results suggest that ferries, or other commercial shipping routes, represent a promising sampling platform to alleviate constraints on access to pelagic environments that currently limit marine biodiversity surveys.

### Differences between ferry routes and subnetworks

Environmental variability and species composition were compared between individual ferry routes to understand which ferry routes were important when building a subnetwork. The routes between Salerno-Ashdod and Venice-Patras were the only two routes that covered the extremities of environmental variability and so were required in any ferry subnetwork to achieve full coverage of the environmental parameter space. Previous research suggests that as long as a biased sampling frame covers the environmental variability present in the whole sampling frame, it does not matter if it is geographically biased (Kadmon et al., 2004; Tessarolo et al., 2014). However, our results demonstrate that the environment subnetwork was not able to accurately predict community composition despite covering environmental variability, and in fact performed similarly to the deliberately biased subnetwork (Figure 3). This highlights that environmental variability is not the only factor affecting biases related to restricted sampling frames. The NMDS analysis showed that the routes covering Salerno-Ashdod and Venice-Patras do not cluster with any other routes suggesting that different species compositions occur on these routes (Supplementary Information). Meanwhile, the community subset, which covered both community composition and environmental variability, predicted species richness in the ‘perfection knowledge’ SSDM most accurately. This result highlights that community composition as well as environmental variability must be considered when selecting ferry routes to be representative sampling frames.

The original ferry route network had a high density of shipping routes in the northwest Mediterranean, coinciding with the region with most biodiversity data available, which we expected to bias the predictive capacity of the sampling SSDMs using this network (Figure 2). However, the community subnetwork, with six routes removed from the northwest basin, was still able to accurately predict community composition suggesting this was not a driving factor in the effectiveness of the ferry routes as a sampling frame (Figure 3). A limitation of using existing community composition knowledge to select ferry routes for sampling is that it requires reliable occurrence data to model a ‘perfect knowledge’ SSDM. Here, the NMDS analysis shows that community composition along the ferry routes is related to the geographical location of the routes, with routes closer together having more similar community composition (Supplementary Information). We also demonstrated that increasing the number of routes within the network and having fewer sampling points along more routes, will lead to improved predictions of community composition. Therefore, we would recommend implementing a large number of ferry routes, at least 8, that cover as many different regions of a study area as possible if pre-exiting occurrence data is unreliable or limited.

### Random and systematic biases in data collection

Reports identifying taxonomic biases in biodiversity surveys are pervasive in the literature, but little is known how taxonomic biases can affect downstream analyses such as species distribution modelling or spatial planning (Donaldson et al., 2016; Di Marco et al., 2017; Troudet et al., 2017). Instead, efforts to reduce bias in species distribution models have largely been directed at spatial and temporal biases in data collection (Kramer-Schadt et al., 2013; Beck et al., 2014; Inman et al., 2021). We demonstrate that different taxa have varying species richness gradients, thus removing different taxonomic groups affected which species richness gradients were revealed. The classes Actinopterygii (fishes) and Chondrichthyes (sharks and rays) both showed highest species richness closest to shore whereas marine mammals were more prevalent offshore. This is unsurprising as Actinopterygii and Chondrichthyes are more closely related, and are largely ectothermic so more constrained by temperature requirements than marine mammals (Losos, 2008; Grady et al., 2019). However, this may have been exaggerated by more marine mammal data being collected offshore from ferries compared to Actinopterygii and Chondrichthyes data which is largely collected by coastal fisheries (Aïssi et al., 2015; Mancusi et al., 2020). As more species in the ‘perfect knowledge’ SSDM belonged to classes Actinopterygii and Chondrichthyes, this meant that SSDMs with marine mammals removed were more similar to the ‘perfect knowledge’ SSDM (Figure 5). This highlights that the proportion of species representing each class has an important influence on the overall species richness gradients captured. If biases lead to certain taxonomic groups being underrepresented then it is unlikely that their species richness gradients would be adequately captured, unless they follow similar distributions to another taxa. To utilise novel methods for biodiversity data collection most effectively, it is important to understand the effect taxonomic bias can have, and how new methods can best reduce current biases.

### Conclusion/Future research

Our study demonstrates that high quality biodiversity data can be collected from biased sampling frames, providing they cover wide areas and diversified habitats. Utilising these biased sampling frames, such as ferries, allows data collection from challenging and remote areas which are often inaccessible to researchers due to logistical and financial constraints. This is particularly relevant for upscaling sampling for emerging biodiversity monitoring techniques, such as eDNA sampling, to reduce current spatial, temporal and taxonomic biases (Pawlowski et al., 2020). This study focused on the ferry routes in the Mediterranean to carry out simulated sampling strategies, but sampling from ferry routes, as well as other commercial vessels, could be carried out across the global shipping network. The efficiency for ferry routes as sampling platforms will depend on the concentration of ferry routes in the study area or region of interest. Global cargo routes are largely concentrated in the North Atlantic, North Pacific and Indian Oceans, linking Europe, North America, East and Southeast Asia. High traffic routes crossing the South Atlantic and South Pacific also connect with Southern Africa, South America and Australasia. These represent key areas where commercial vessels could contribute to closing gaps in biodiversity data (Wang, C. and Wang, 2011). These areas also coincide with those most affected by human impacts emphasising the need for regular monitoring to understand effects on biodiversity (Halpern et al., 2008; Pirotta et al., 2019). The workflow presented here can be used as a template to evaluate the efficiency of a shipping route network in a study area of interest before undertaking sampling. This study focused on the impact of sampling strategies on species distribution models, which are frequently used as conservation features in marine spatial planning to designate protected areas. Therefore, our findings confirm that biased sampling, if designed adequately, can provide a useful data basis for marine species and management of marine environments.

