## Supplementary Information for "Sampling from commercial vessel routes can capture marine biodiversity distributions effectively"

| | |
|---|---|
| **Supplementary Table S1** | Marine predator species used in ‘perfect knowledge’ stacked-species distribution model |
| **Supplementary Methods S1** | Principal component analysis to build environmental variability map |
| **Supplementary Table S2** | Evaluation metrics for ‘perfect knowledge’ stacked-species distribution model |
| **Supplementary Table S3** | Importance of the environmental predictors in the ‘perfect knowledge’ stacked-species distribution model |
| **Supplementary Table S4** | ANOVA tables for stacked-species distribution model comparisons between sampling strategies and ferry route subnetworks |
| **Supplementary Table S5** | Individual ferry route and ferry subnetwork’s length, species covered and climatic bias score |
| **Supplementary Figure S1** | Map of Mediterranean highlighting which regions are covered by ferry routes and NMDS plot showing community composition differences among ferry routes. |

**Supplementary Table S1**

List of 43 marine predator species that had > 40 occurrence points from combined data from online repositories, GBIF, OBIS and EurOBIS, Accobams and the Medlem database. Total length and tropic level as reported by FishBase (Actinopterygii and Chondrichthyes) or SeaLifeBase (Mammalia and Reptilia). Marine predator defined as having total length ≥ 1m and trophic level ≥ 4. Species that do not meet these criteria but were retained are denoted with an Asterix.

| Species | Common name | Class | Total length (cm) | Trophic Level | Number of occurrences |
|---|---|---|---|---|---|
| *Conger conger* | European Conger | Actinopterygii | 573.3 | 4.3 | 649 |
| *Dentex dentex* | Common dentex | Actinopterygii | 100 | 4.5 | 372 |
| *Echelus myrus* | Painted eel | Actinopterygii | 100 | 4.3 | 45 |
| *Epinephelus aeneus* | White grouper | Actinopterygii | 120 | 4 | 49 |
| *Epinephelus marginatus* | Dusky grouper | Actinopterygii | 150 | 4.4 | 969 |
| *Fistularia commersonii* | Bluespotted cornetfish | Actinopterygii | 160 | 4.3 | 63 |
| *Lophius piscatorius* | Angler | Actinopterygii | 200 (SL) | 4.5 | 602 |
| *Merluccius merluccius* | European hake | Actinopterygii | 140 | 4.4 | 1179 |
| *Mola mola* | Ocean sunfish | Actinopterygii | 333 | 3.3 | 3530 |
| *Molva dypterygia* | Blue ling | Actinopterygii | 155 | 4.5 | 119 |
| *Muraena helena* | Mediterranean moray | Actinopterygii | 150 | 4.2 | 1061 |
| *Ophisurus serpens* | Serpent eel | Actinopterygii | 250 | 4.1 | 47 |
| *Pomatomus saltatrix* | Bluefish | Actinopterygii | 130 | 4.5 | 82 |
| *Seriola dumerili* | Greater amberjack | Actinopterygii | 190 | 4.5 | 118 |
| *Sphyraena sphyraena* | European barracuda | Actinopterygii | 165 | 4 | 159 |
| *Sphyraena viridensis* | Yellowmouth barracuda | Actinopterygii | 128 (FL) | 4.3 | 54 |
| *Thunnus alalunga* | Albacore | Actinopterygii | 140 (FL) | 4.3 | 270 |
| *Thunnus thynnus* | Bluefin tuna | Actinopterygii | 458 | 4.5 | 177 |
| *Xiphias gladius* | Swordfish | Actinopterygii | 455 | 4.5 | 48 |
| *Zu cristatus* | Scalloped ribbonfish | Actinopterygii | 118 (SL) | 4.5 | 204 |
| *Alopias vulpinas* | Common thresher | Chondrichthyes | 573.3 | 4.5 | 114 |
| *Carcharhinus longimanus* | Oceanic whitetip shark | Chondrichthyes | 400 | 4.2 | 77 |
| *Cetorhinus maximus** | Basking shark | Chondrichthyes | 1520 | 3.2 | 142 |
| *Dasyatis pastinaca** | Common stingray | Chondrichthyes | 64 (WD) | 4.1 | 163 |
| *Echinorhinus brucus* | Bramble shark | Chondrichthyes | 310 | 4.4 | 41 |
| *Hexanchus griseus* | Bluntnose sixgill shark | Chondrichthyes | 482 | 4.5 | 140 |

| | | | | | |
|---|---|---|---|---|---|
| *Isurus oxyrinchus* | Short-fin mako shark | Chondrichthyes | 445 | 4.5 | 81 |
| *Mobula mobular** | Giant devil ray | Chondrichthyes | 520 (WD) | 3.7 | 874 |
| *Myliobatis aquila** | Common eagle ray | Chondrichthyes | 183 (WD) | 3.6 | 119 |
| *Prionace glauca* | Blue shark | Chondrichthyes | 400 | 4.4 | 324 |
| *Raja clavata** | Thornback ray | Chondrichthyes | 139 | 3.8 | 431 |
| *Squalus acanthias** | Spiny dogfish | Chondrichthyes | 95 | 4.4 | 78 |
| *Torpedo marmorata* | Marbled electric ray | Chondrichthyes | 100 | 4.5 | 116 |
| *Balaenoptera physalus* | Fin whale | Mammalia | 2700 | 3.2-4.3 | 1245 |
| *Delphinus delphis* | Short-beaked common dolphin | Mammalia | 260 | 4.5 | 1693 |
| *Globicephala melas* | Long-finned pilot whale | Mammalia | 670 | 4.5 | 1147 |
| *Grampus griseus* | Risso’s dolphin | Mammalia | 380 | 4.36-4.54 | 410 |
| *Orcinus orca* | Killer whale | Mammalia | 980 | 4.5-4.6 | 115 |
| *Physeter macrocephalus* | Sperm whale | Mammalia | 2400 | 4.5-4.7 | 2307 |
| *Stenella coeruleoalba* | Striped dolphin | Mammalia | 260 | 4.5 | 7822 |
| *Tursiops truncatus* | Bottlenose dolphin | Mammalia | 380 | 4.5 | 3991 |
| *Ziphius cavirostris* | Cuvier’s beaked whale | Mammalia | 750 | 4.5 | 113 |
| *Caretta caretta** | Loggerhead turtle | Reptilia | 125 (CL) | 3.5-3.6 | 1557 |

Ray species *D. pastinaca* and *Torpedo torpedo* were retained due to having trophic levels greater than 4. Total length is not reported for these species so retained despite their width being less than 1 metre.

*R. clavata* and *S. acanthias* were retained as they were so close to threshold.

*Mola mola, Cethorhinus maximus, Mobular mobular, Myliobatis aquila* and *Caretta caretta* were retained as they are classed as marine megafauna species which can exert top down effects on ecosystems, similar to that of predators (Pimiento et al., 2020).

Pimiento, C., Leprieur, F., Silvestro, D., Lefcheck, J., Albouy, C., Rasher, D., Davis, M., Svenning, J.-C. and Griffin, J. 2020. Functional diversity of marine megafauna in the Anthropocene. *Science Advances.* **6**(16), peaay7650.

**Supplementary Methods S1**

Principal component analysis (PCA) was used to create an environmental variability map. Initially, PCA was conducted on a correlation matrix of standardized environmental predictors used to create the SSDM using the prcomp() function in the 'stats' R package. The first four principal components were retained for downstream analysis as they explained >80 % of the variability in the environmental predictors. Site scores, i.e. weighted linear combinations of the environmental predictors, were used to produce surface maps of each of the first four principal components to visualize the main gradients of environmental variability in the study area. The principal components were collapsed into one surface map of environmental variability by summing the site scores of each principal component weighted according to its contribution as following the equation:

$$EV = (0.3665*PC1) + inv(0.2435*PC2) + (0.164*PC3) + (0.1121*PC4)$$

Mean bathymetry made an important contribution to PC1 and PC2 (factor loading >0.4) but a high score in PC1 related to shallow bathymetry while a high score in PC2 related to greater depths. Therefore, PC2 was inverted otherwise the site scores offset each other and the variability was lost. The final 'environmental variability map' is unitless and shows the main gradients in environmental variability in the study area.

The first principal component explained 36.65 % of the variability in the environmental predictors where the highest values correspond to shallow bathymetry and low sea surface temperatures but high values of chlorophyll concentration and sea surface temperature range. The second principal component explained 24.35 % of the variability and represents variability related to distance from shore, where bathymetry is deepest further from shore. The third principal component explained 16.4 % of the variation where larger values correlated to the largest bathymetric slope. The fourth principal component explained 11.21 % of the variation with the northern Adriatic clearly being the most different area due to having a high chlorophyll concentration and lower sea surface temperature.

The weighted overlay of the principal components shows the overall trends in environmental variability where the values are unitless but the larger the range between values represents areas with the most different environmental conditions. There are two main trends in environmental variability, 1) a difference between the north-western and south-eastern basin, 2) a gradient with distance from shore. The northern tip of the Adriatic Sea is clearly most different from the rest of the Mediterranean.

Principal components respective contribution ratios.

| | PC1 | PC2 | PC3 | PC4 | PC5 | PC6 |
|---|---|---|---|---|---|---|
| **Eigenvalue** | 1.4828 | 1.2088 | 0.992 | 0.8203 | 0.65213 | 0.50760 |
| **Contribution ratio (%)** | 0.3665 | 0.2435 | 0.164 | 0.1121 | 0.07088 | 0.04294 |
| **Cumulative contribution (%)** | 0.3665 | 0.6100 | 0.774 | 0.8862 | 0.95706 | 1.0000 |

Eigenvectors. Factor loadings >0.4 are highlighted in bold.

| Variable (units) | PC1 | PC2 | PC3 | PC4 |
|---|---|---|---|---|
| **Mean bathymetry (m)** | **-0.4281** | **0.5227** | -0.2578 | 0.0677 |
| **Mean sea surface temperature (°C)** | **-0.4773** | -0.3344 | 0.3005 | 0.2529 |
| **Mean chlorophyll concentration (mg/m$^3$)** | **0.4436** | 0.1910 | -0.0693 | **0.8660** |
| **Sea surface temperature range (°C)** | **0.4783** | 0.3533 | -0.1049 | **-0.4166** |
| **Bathymetric slope (°)** | -0.1490 | -0.2761 | **-0.9079** | 0.0511 |
| **Distance to shore (km)** | -0.3756 | **0.6143** | 0.0568 | 0.0735 |

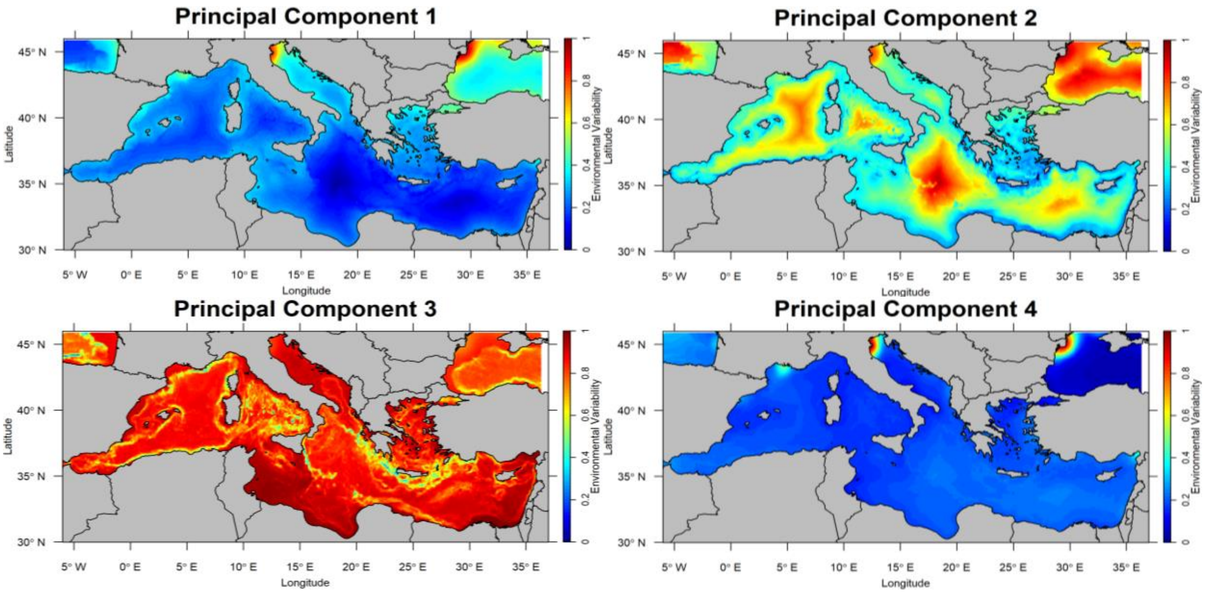


Principal component scores projected onto the study area.

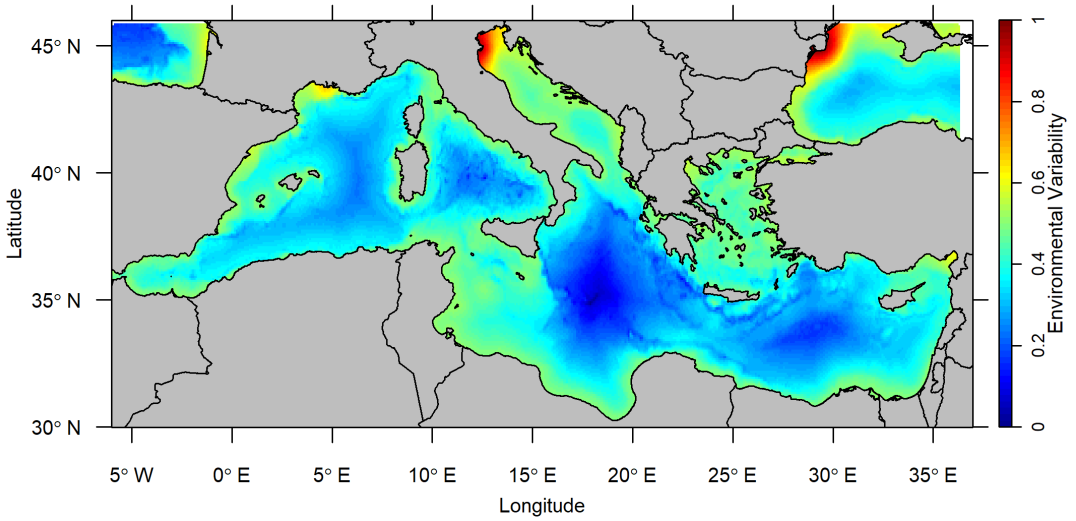


Environmental variability map using weighted overlay of the first four principal components.

**Supplementary Table S2**

Results from six metrics used to evaluate the prediction accuracy of species assemblage predictions in the 'perfect knowledge' SSDM.

| | Species Richness Error | Prediction Success | Kappa | Specificity | Sensitivity | Jaccard |
|---|---|---|---|---|---|---|
| Mean | 19.06 | 5.98 | 0.995029 | 0.544447 | 0.983425 | 0.065596 |
| SD | 7.229589 | 1.797052 | 0.000441 | 0.172317 | 0.124641 | 0.059542 |

**Supplementary Table S3**

Pearson correlation coefficient calculated from the difference between a full model and one with each environmental variable omitted in turn for individual species models then averaged across species.

| | Mean bathymetry | Mean sea surface temperature | Mean chlorophyll concentration | Mean temperature range | Bathymetric slope | Distance from shore |
|---|---|---|---|---|---|---|
| Mean | 14.59 | 31.76 | 13.17 | 16.7 | 8.31 | 15.47 |
| SD | 6.56 | 13.74 | 5.1 | 5.53 | 1.96 | 8.32 |

**Supplementary Table S4**

**Analysis of Variance Tables**

One-way ANOVA to evaluate the impact of different ferry route subnetworks on correlation coefficients.

| Factor | Df | Sum Sq | Mean Sq | F value | P value |
|---|---|---|---|---|---|
| Sampling frame | 2 | 1.20122 | 0.60061 | 342.96 | <0.001* |

Tukey's Post Hoc Test to evaluate which ferry route subnetworks differed from each other.

| Group 1 | Group 2 | Estimate | Conf. low | Conf. high | P value |
|---|---|---|---|---|---|
| Biased | Community | 0.384 | 0.336 | 0.432 | <0.001* |
| Biased | Environment | -0.0952 | -0.142 | -0.0487 | <0.001* |
| Community | Environment | -0.479 | -0.527 | -0.431 | <0.001* |

One-way ANOVA to evaluate the effect of the number of ferry routes in a ferry subnetwork. Dependent variable was square transformed prior to analysis.

| Factor | Df | Sum Sq | Mean Sq | F value | P value |
|---|---|---|---|---|---|
| Number of ferries | 5 | 1.2987 | 0.259745 | 15.286 | <0.001* |

Tukey's Post Hoc Test to evaluate which ferry subnetworks differed depending on the number of ferry routes.

| Group 1 | Group 2 | Estimate | Conf. low | Conf. high | P Value |
|---|---|---|---|---|---|
| 2 | 4 | 0.116 | -0.0558 | 0.289 | ns |
| 2 | 6 | 0.149 | -0.0233 | 0.321 | ns |
| 2 | 8 | 0.379 | 0.207 | 0.552 | <0.001* |
| 2 | 10 | 0.364 | 0.191 | 0.536 | <0.001* |
| 2 | 12 | 0.362 | 0.189 | 0.534 | <0.001* |
| 4 | 6 | 0.0325 | -0.140 | 0.205 | ns |
| 4 | 8 | 0.263 | 0.0907 | 0.435 | <0.001* |
| 4 | 10 | 0.247 | 0.0750 | 0.419 | <0.001* |
| 4 | 12 | 0.245 | 0.0730 | 0.418 | <0.001* |
| 6 | 8 | 0.230 | 0.0582 | 0.403 | <0.001* |
| 6 | 10 | 0.215 | 0.0425 | 0.387 | <0.001* |
| 6 | 12 | 0.213 | 0.0405 | 0.385 | <0.001* |
| 8 | 10 | -0.0157 | -0.188 | 0.157 | ns |
| 8 | 12 | -0.0177 | -0.190 | 0.155 | ns |
| 10 | 12 | -0.00194 | -0.174 | 0.170 | ns |

Three-way ANOVA to evaluate the impact of removing specific taxa from stacked species distribution models across sampling strategies (random vs regular) and sampling sizes (25, 50, 100 sampling points). Dependent variable (the correlation coefficient) was square transformed prior to analysis.

| Factor | Df | Sum Sq | Mean Sq | F value | P value |
|---|---|---|---|---|---|
| Strategy | 1 | 0.08385 | 0.083854 | 7.9500 | <0.05* |
| Size | 2 | 0.26295 | 0.131475 | 12.4648 | <0.001* |
| Taxa removed | 2 | 0.18388 | 0.091941 | 8.7167 | <0.001* |

Tukey’s Post Hoc Test to look at pairwise differences between taxa removed, sampling strategy and sampling size.

| Term | Group 1 | Group 2 | Estimate | Conf. low | Conf. high | P value |
|---|---|---|---|---|---|---|
| Strategy | Random | Regular | 0.0522 | 0.0155 | 0.0889 | <0.05* |
| Size | 25 | 50 | -0.0400 | -0.0936 | 0.0136 | ns |
| Size | 25 | 100 | 0.0715 | 0.0170 | 0.126 | <0.05* |
| Size | 50 | 100 | 0.112 | 0.0580 | 0.165 | <0.001* |
| Taxa removed | Chondrichthyes | Mammalia | 0.106 | 0.0428 | 0.169 | <0.001* |
| Taxa removed | Chondrichthyes | Random | 0.0299 | -0.0241 | 0.0840 | ns |
| Taxa removed | Mammalia | Random | -0.0758 | -0.130 | -0.0217 | <0.05* |

**Supplementary Table S5**

The ferry route or ferry route subnetwork length, the number of species distributions overlapping with the ferry route or subnetwork, and the climatic bias index.

| Ferry Route | Ferry length (no. grid cells covered) | Number of species | Climatic Bias Index |
|---|---|---|---|
| SaAs | 404 | 36 | 0.3944031 |
| TaGe | 278 | 41 | 0.17952261 |
| VaSa | 218 | 36 | 0.42306601 |
| VePa | 194 | 38 | 1.0530864 |
| CiBa | 144 | 39 | 0.38805581 |
| AtIz | 81 | 34 | 1.1377996 |
| ToAl | 69 | 39 | 0.6424208 |
| ToBas | 55 | 38 | 0.43982341 |
| KaMi | 52 | 36 | 1.1377996 |
| ToAj | 48 | 39 | 0.77694461 |
| NiBas | 43 | 39 | 0.1230776 |
| LiGA | 42 | 40 | 1.1377996 |
| ToIR | 42 | 39 | 0.60432561 |
| SaBas | 35 | 38 | 0.39494239 |
| NiIR | 32 | 37 | 0.797778 |
| All ferries | 1744 | 42 | 0.06914759 |
| Biased subnetwork | 513 | 41 | 0.3147127 |
| Community subnetwork | 1462 | 42 | 0.07962026 |
| Environment subnetwork | 598 | 39 | 0.08448943 |

**Supplementary Figure S1**

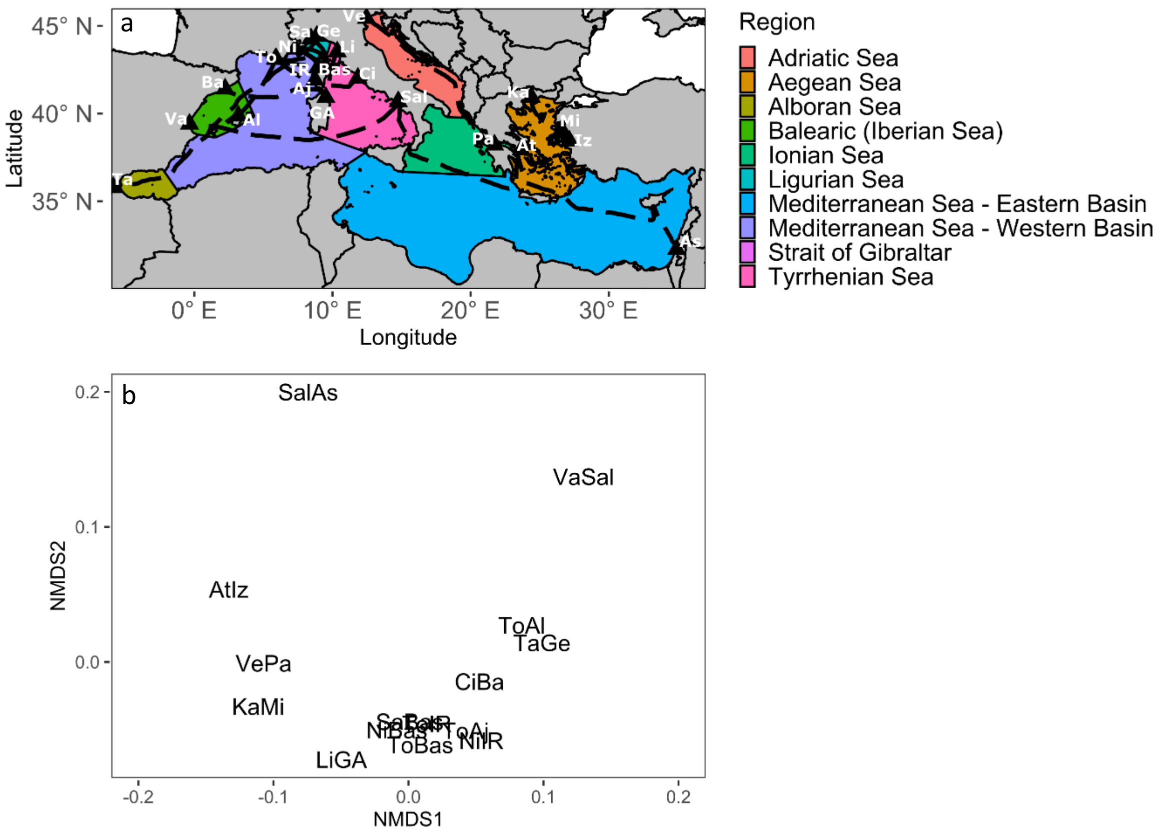


a) Map of the Mediterranean showing regions covered by different ferry routes. b) Non-metric multidimensional scaling plot based on Bray-Curtis dissimilarity matrix for species composition between different ferry routes.
